# Baculovirus-mediated gene transfer enables functional expression of the PIEZO1 ion channel in isolated muscle satellite cells

**DOI:** 10.1101/2025.09.01.673443

**Authors:** Akira Murakami, Ayuno Masuda, Takumi Hirakawa, Kotaro Hirano, Masaki Tsuchiya, Yusuke Ono, Takashi Murayama, Yuji Hara

## Abstract

Primary tissue stem cells are useful not only for basic cell biological research but also for therapeutic applications: however, their broader utility is often limited by technical challenges, such as a low efficiency of exogenous gene expression. PIEZO1 is a large mechanosensitive ion channel that plays an important role in muscle-resident stem cells, known as muscle satellite cells (MuSCs), during muscle regeneration. In this study, we developed a method for the ectopic expression of PIEZO1 in isolated MuSCs. Using a baculovirus vector system, we expressed PIEZO1 in myoblast C2C12 cells. Following optimization of the infection condition, we achieved robust PIEZO1 expression in isolated MuSCs during activated and differentiated states, with appropriate subcellular localization and ion channel activity. Importantly, the baculovirus-mediated PIEZO1 expression restored the reduced proliferative capacity of *Piezo1*-deficient MuSCs to a level comparable to wild-type cells, indicating that the exogenously expressed PIEZO1 is functionally equivalent to the endogenous protein. Overall, we established an efficient method for the transfer of the *Piezo1* gene into isolated MuSCs, which should provide a versatile platform to study other large proteins in MuSCs.

**Summary Statement:** A baculovirus-based method was developed, enabling robust and functional PIEZO1 expression in isolated muscle satellite cells and providing a platform to study large proteins in isolated stem cells.

## Introduction

Tissue-resident stem cells have an essential role in maintaining the homeostasis of adult tissues (Simons and Clevers, 2011). Under physiological conditions, these cells generally exist in a quiescent state but are activated in response to environmental stimuli, upon which they enter the cell cycle, proliferate, and differentiate (Urbán and Cheung, 2021). Isolated tissue stem cells, particularly those maintained in a quiescent state, serve not only as *ex vivo* models for examining mechanisms of tissue regeneration, but are also promising candidates for transplantation-based therapies targeting tissues (Zakrzewski et al., 2019). To support these applications, strategies for introducing and overexpressing genes of interest in isolated tissue stem cells have been evaluated (Hamann et al., 2019): however, achieving efficient gene transfer, particularly in quiescent stem cells, remains a challenge (Arnett et al., 2014).

Muscle satellite cells (MuSCs) are skeletal muscle-resident stem cells that are indispensable for myofiber regeneration (Relaix and Zammit, 2012). Following injury, they are activated and differentiate into myoblasts, which fuse with damaged myofibers or other myoblasts to generate new myofibers. During this process, MuSCs express distinct transcription factors at each stage: PAX7 in quiescence, MYOD during activation, and MYOG upon differentiation (Relaix et al., 2021). Primary MuSCs isolated from skeletal muscle recapitulate these phenotypical and transcriptional profiles *ex vivo*. MuSCs proliferate when cultured in growth medium and differentiate in serum-starved medium to induce myoblast fusion and myotube formation (Danoviz and Yablonka-Reuveni, 2011). Thus, these cells are used as a robust model for analyzing MuSC-dependent muscle regeneration.

Previously, we demonstrated that force-sensing proteins play an important role in MuSC function during myofiber regeneration (Hirano et al., 2025; Hirano et al., 2023). PIEZO1 is a mechanosensitive ion channel activated by membrane tension (Ge et al., 2015). Studies using tissue-specific *Piezo1* knockout mice revealed that PIEZO1 acts as a mechanosensor in various tissues, including neural progenitor cells (Pathak et al., 2014), chondrocytes (Lee et al., 2014), and blood and lymphatic vessels (Li et al., 2014; Nonomura et al., 2018). We found that *Piezo1* is highly expressed in MuSCs, and generated MuSC-specific *Piezo1*-deficient mice (*Piezo1^flox/flox^*; *Pax7^CreERT2^*^/+^, hereafter referred to as *Piezo1*-cKO mice) to examine PIEZO1 function (Hirano et al., 2023). These mice showed delayed muscle regeneration following injury, at least in part, because of impaired proliferation and mitotic catastrophe, both of which resulted from defects in Rho-mediated signaling. Moreover, using *Piezo1*-tdTomato mice, in which the fluorescent protein tdTomato was fused to the C-terminus of endogenous PIEZO1 protein, we demonstrated that PIEZO1 is localized to the midbody during cytokinesis. Although these results have clarified the role of PIEZO1 in MuSCs, further studies using ectopic PIEZO1 expression in isolated MuSCs are needed to elucidate PIEZO1 functions.

Viral vectors are an attractive solution for gene transfer into primary stem cells (Bulcha et al., 2021): however, because PIEZO1 is encoded by a ∼7.7 kb gene, its size was expected to be exceeds packaging capacity of viral vectors, particularly when fused to other genes such as *Gfp* (Sweeney and Vink, 2021). For example, while lentiviral vectors were successfully applied to various isolated stem cells, including hematopoietic stem cells for *ex vivo* gene therapy (Aiuti et al., 2013), their low packaging capacity is likely insufficient for delivering the *Piezo1* gene. In contrast, baculoviral vectors have a much higher packaging capacity and can deliver genes up to 38 kb into a mammalian cell line (Cheshenko et al., 2001). Baculovirus-mediated gene transfer has also been used in human embryonic stem cells (Zeng et al., 2007), induced pluripotent stem cells (Mansouri et al., 2016), and various muscle-related cells, including primary rat myoblasts (Shen et al., 2008) and primary trout MuSCs (Jackson et al., 2013): however, these studies have only been validated in the delivery of small reporter genes such as *egfp*, and there have been no reports demonstrating the expression of large genes like *Piezo1* in isolated tissue stem cells.

In this study, we developed a method for the exogenous expression of the *Piezo1* gene in MuSCs isolated from mouse skeletal muscle. Because of the large size of the *Piezo1* gene, a conventional transfection technique (e.g., lipofection) was inefficient at expressing the PIEZO1 protein even in the murine myoblast cell line C2C12. Therefore, we used baculovirus-mediated gene transfer and optimized the infection conditions. As a result, we achieved a high expression efficiency of *Piezo1* in isolated MuSCs, despite its larger size.

## Results

### Baculovirus-mediated gene transfer enables the expression of a large gene *Piezo1* in C2C12 cells

The murine myoblast cell line C2C12 was used as a model to establish an effective method for exogenously expressing the *Piezo1* gene. First, we evaluated the lipofection method for transferring the *Piezo1* gene fused to the *egfp* gene. Transfection efficiency was assessed by fluorescence-activated cell sorting (FACS), which quantified EGFP signal intensity at the single-cell level. C2C12 cells transfected with a plasmid encoding *egfp*-fused *Piezo1* displayed a fluorescence profile similar to that of the control cells, indicating that the *Piezo1*-expressing plasmid was inefficiently introduced into the cells. In contrast, cells transfected with a plasmid encoding only *egfp* showed a subpopulation with a higher fluorescence intensity compared with that in the control group (Supplementary Fig. 1A). The mean fluorescence intensity of the *egfp*-only group was higher compared with that of the *Piezo1-egfp* and control groups (Supplementary Fig. 1B). Thus, the results indicated that lipofection is not suitable for introducing *Piezo1* into C2C12 cells, likely because of its large size.

As an alternative approach, we adopted a baculovirus vector system to transfer the large *Piezo1* gene into C2C12 cells. A donor plasmid was constructed to express the vesicular stomatitis virus G-protein (VSVG) under the control of the insect cell-specific polyhedrin promoter. It was also designed to express PIEZO1 fused with EGFP (PIEZO1-EGFP) at the C-terminus under the control of the CMV promoter (Fig. 1A) (Hofmann et al., 1995). Pseudotyped baculoviruses expressing the VSVG envelope protein exhibit increased infectivity in mammalian cells (Barsoum et al., 1997). The donor plasmids harboring *Piezo1* (pFB-PIEZO1) or the empty vector (pFB) were transformed into *E. coli* DH10Bac containing bacmid, a modified baculoviral genome (Airenne et al., 2003). In DH10Bac, Tn7 transposon-mediated transposition resulted in the insertion of the target sequence into the bacmid, which was then transfected into Sf9 insect cells. Baculovirus envelope protein GP64 expression in Sf9 cells confirmed the production of recombinant baculoviruses from the pFB and pFB-PIEZO1 constructs (Supplementary Fig. 2) (Kitts and Green, 1999).

**Figure 1.**
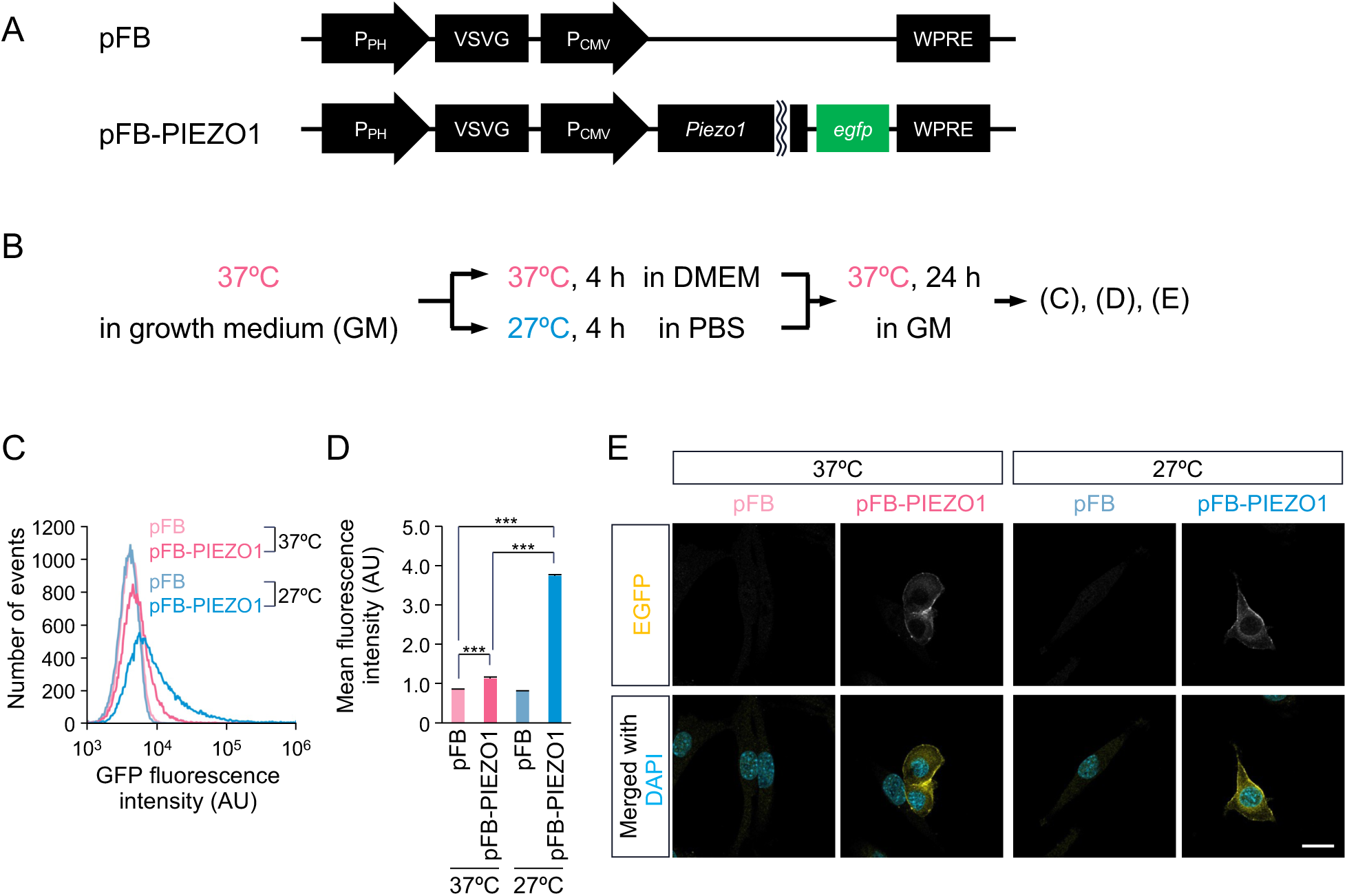
Baculovirus-mediated expression of PIEZO1-EGFP in C2C12 cells. (A) A schematic representation showing a recombinant donor plasmid expressing PIEZO1-EGFP (pFB-PIEZO1). The plasmid lacking the PIEZO1-EGFP sequence (pFB) was used as a control. PH; polyhedrin, VSVG; vesicular stomatitis virus G-protein, WPRE; woodchuck hepatitis post-transcriptional regulatory element. (B) Overview of baculovirus-mediated gene transfer into C2C12 cells. (C) Flow cytometry analyses of the GFP fluorescence intensity in C2C12 cells (30,000 cells). (D) The mean fluorescence intensity among the cells analyzed by the flow cytometry. N = 3, Mean + SE, \*\*\**p* < 0.001. (E) The immunofluorescent staining of PIEZO1 expressed in C2C12 cells. PIEZO1-EGFP was detected using an anti-GFP antibody (yellow). DAPI (blue) indicated the nuclei. Scale bar: 20 μm.

Previous studies have indicated that culture temperature and medium composition during infection affect transduction efficiency (Sung et al., 2014). C2C12 cells were infected with recombinant baculoviruses under the conditions presented in Fig. 1B and cultured for 24 h. FACS analyses revealed that cells infected with pFB-PIEZO1 under all tested conditions exhibited a subpopulation with higher EGFP fluorescence compared with the control groups. Notably, infection at 27°C in PBS yielded the highest proportion of EGFP-positive cells (Fig. 1C). Consistently, the mean fluorescence intensity of this subpopulation was higher compared with that of the other groups (Fig. 1D). To determine the underlying mechanism, we compared PIEZO1 expression by changing both the medium (GM or PBS) and temperature (37°C or 27°C) (Supplementary Fig. 3A). The results indicated that infection at 27°C in PBS yielded the highest expression of PIEZO1 in C2C12 cells (Supplementary Fig. 3B). Immunofluorescence staining revealed EGFP signals only in cells infected with pFB-PIEZO1, whereas no significant difference in signal intensity was observed between cells infected at 37°C in DMEM or at 27°C in PBS (Fig. 1E). Taken together, these results indicate that the baculovirus vector system enables the successful transfer of the large *Piezo1* gene into C2C12 cells. Moreover, infection at 27°C in PBS enhances the proportion of transduced cells, rather than increasing the expression level per cell, compared with infection at 37°C in DMEM.

### PIEZO1 expressed via baculovirus in C2C12 cells is functional as an ion channel

The functional properties of the PIEZO1 ion channel ectopically expressed in C2C12 cells were evaluated. The molecular weight of the PIEZO1-EGFP is approximately 300 kDa based on its amino acid sequence. An immunoblot analysis using an anti-GFP antibody revealed a band of approximately 300 kDa in cells infected with pFB-PIEZO1, suggesting that the full-length PIEZO1-EGFP protein was successfully expressed (Fig. 2A). Immunofluorescence staining revealed that EGFP signals localized at the cell periphery. In a line-scan analysis, EGFP fluorescence displayed a pattern opposite to nuclei staining (DAPI), indicating that PIEZO1-EGFP was predominantly localized to the plasma membrane (Fig. 2B). To determine whether exogenously expressed PIEZO1-EGFP shows ion channel activity, calcium imaging was performed using the ratiometric Ca^2+^ indicator Fura-2 AM. PIEZO1-deficient C2C12 cells, previously established in our laboratory (Tsuchiya et al., 2018), were infected with either pFB-PIEZO1 or the control. Upon stimulation with Yoda1, a PIEZO1-specific agonist, Ca^2+^ influx was observed exclusively in cells infected with pFB-PIEZO1. An increase in the fluorescence ratio (Δratio) was greater in cells infected at 27°C in PBS compared with those infected at 37°C in a standard medium (Fig. 2C): however, the difference was less pronounced than that observed by immunoblotting (Fig. 2A). This suggests that the conditions of 27°C in PBS increased the proportion of transduced cells rather than the expression level per cell, which is consistent with the data shown in Fig. 1E. These results demonstrate that the baculovirus-mediated gene transfer system enables the functional expression of the large PIEZO1 ion channel in the plasma membrane of C2C12 cells.

**Figure 2.**
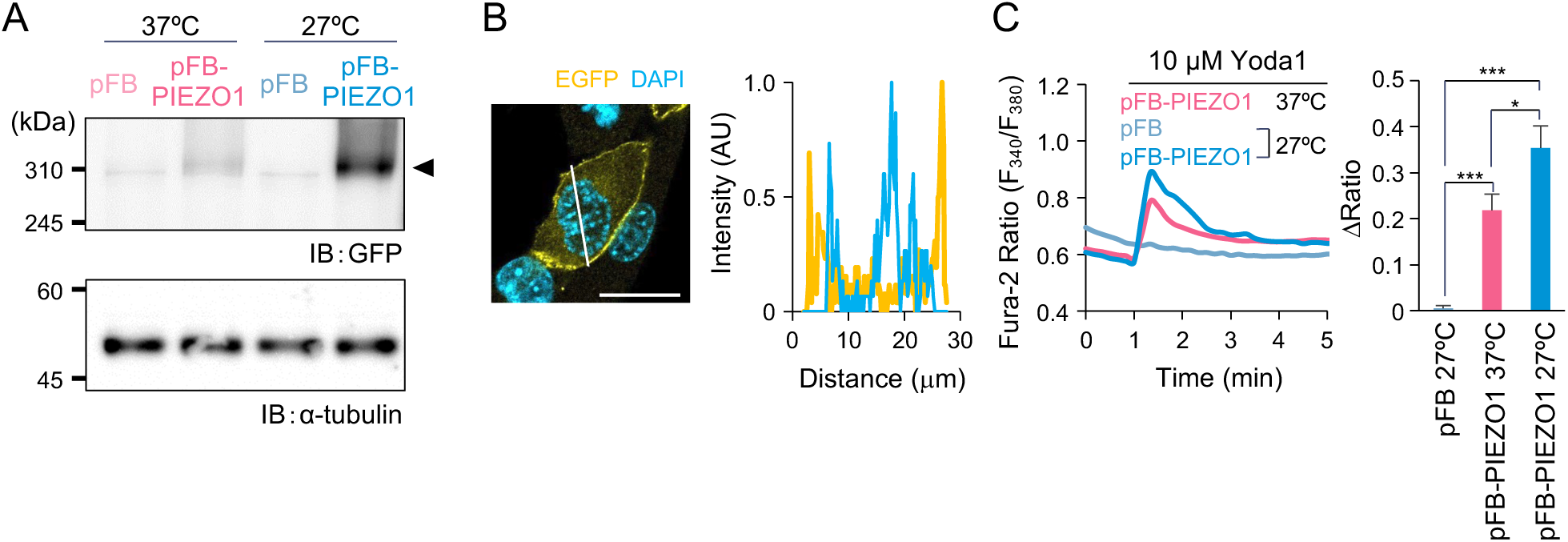
Functional characterization of PIEZO1-EGFP as an ion channel expressed using a baculovirus system in C2C12 cells. (A) A representative image of immunoblot analyses on PIEZO1-EGFP protein expression. After preparation of lysates from C2C12 cells infected with the recombinant baculovirus, protein bands corresponding to PIEZO1 and α-tubulin were detected using an anti-GFP antibody and an anti-α-tubulin antibody, respectively. α-tubulin was used as loading control. (B) A fluorescence intensity profile of PIEZO1-EGFP in C2C12 transduced with recombinant baculovirus. The intensity profile along the white line was shown in the left panel. PIEZO1-EGFP was detected using anti-GFP antibody (yellow). DAPI (blue) indicated the nuclei. Scale bar: 20 μm. (C) Fura-2 ratiometric measurements (F340/F380) of Yoda1-induced Ca^2+^ mobilization in *Piezo1*- deficient C2C12 cells transduced with the recombinant baculovirus under the indicated conditions (left). Changes in Fura-2 ratio (1′Ratio) were quantified in right panel. n = 99, Mean + SE, \**p* < 0.05, \*\*\**p* < 0.001.

### The baculovirus system yields high *Piezo1* gene expression in isolated MuSCs

Next, the *Piezo1* gene transfer conditions optimized in C2C12 cells were applied to MuSCs isolated from mice. As indicated above, MuSCs undergo state transitions during *in vitro* culture, which are reflected in the expression of specific transcription factors (Relaix et al., 2021). For example, PAX7 and MYOD serve as markers for the stem-like state and activated state from quiescence, respectively. Under our experimental conditions, 88% of the MuSCs that were fixed 0.5 h after isolation were PAX7⁺/MYOD⁻, which indicated a quiescent state. At 24 h post-isolation, 72% of the cells expressed PAX7⁺/MYOD⁺, which suggested activation. By 144 h, a population of PAX7⁻/MYOD⁺ cells emerged that represented a transition from MuSCs to myoblasts (Supplementary Fig. 4). Because baculovirus infection efficiency depends on the cell state (Valeri et al., 2024), the MuSCs were infected with baculovirus 0.5, 24, or 144 h after isolation using the 27°C in PBS conditions (Fig. 3A). Immunofluorescent analysis revealed EGFP signals in cells infected with pFB-PIEZO1, but not in those infected with the control at every time point (Fig. 3B). The proportion of EGFP-positive cells was 24% at 0.5 h, 74% at 24 h, and 46% at 144 h following isolation (Fig. 3C). These results indicated that the baculovirus vector system can efficiently transfer *Piezo1* in isolated MuSCs. Furthermore, while the system was applicable to inactive and activated MuSCs, it achieved a higher gene transfer efficiency in activated MuSCs .

**Figure 3.**
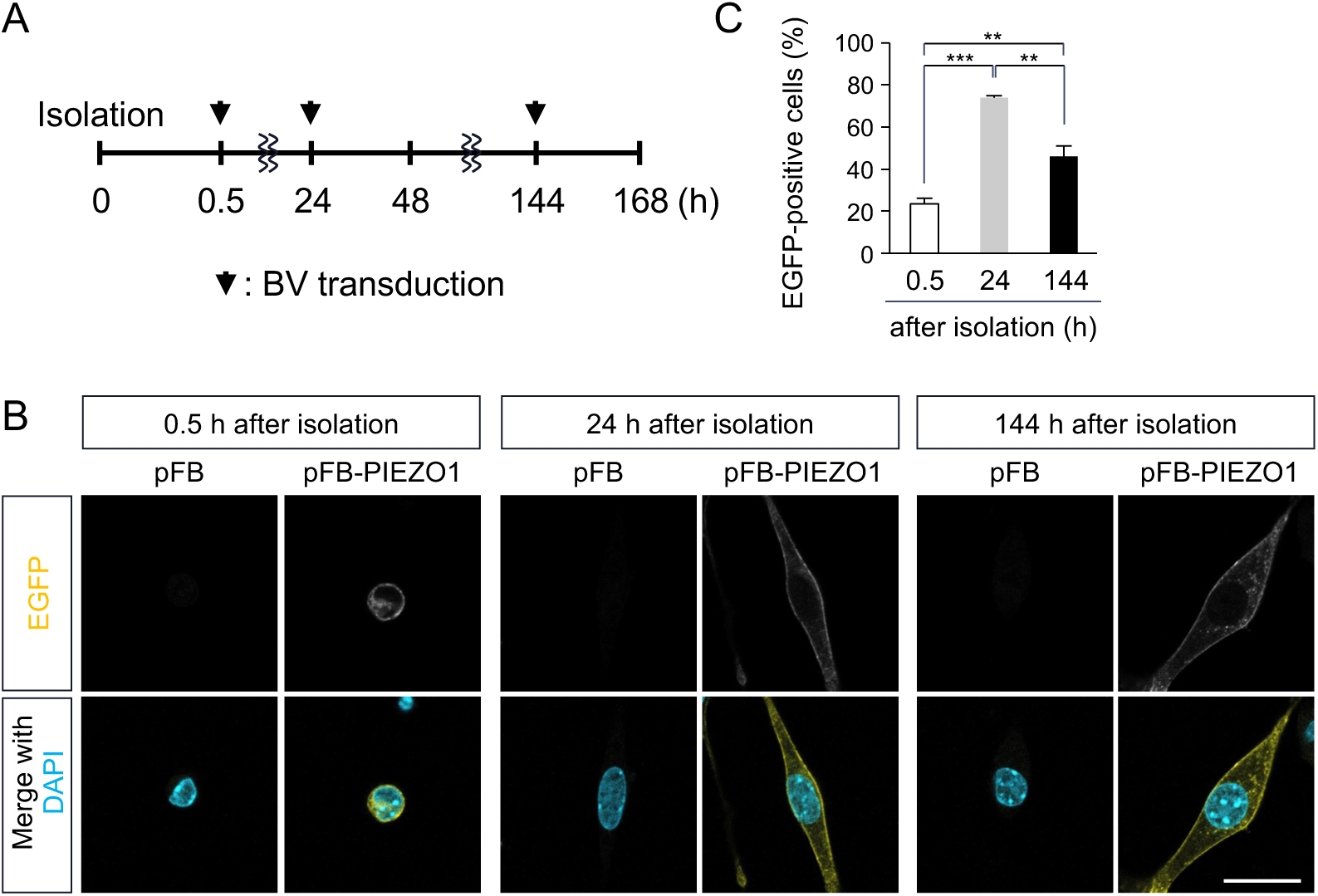
Baculovirus-mediated expression of PIEZO1-EGFP in muscle satellite cells (MuSCs) isolated from mice. (A) Schematic showing baculovirus transduction into MuSCs isolated from mouse limb muscle. (B) The fluorescence immunostaining analysis of PIEZO1-EGFP expression in isolated MuSCs. MuSCs were transduced with recombinant baculovirus at 0.5, 24, and 144 hs after isolation, representing quiescent MuSCs, activated MuSCs, and differentiated myoblasts, respectively. PIEZO1 were detected using an anti-GFP antibody (yellow). DAPI (blue) indicated the nuclei. Scale bar: 30 μm. (C) The percentage of PIEZO1-expressing cells transduced with recombinant baculovirus at 0.5, 24, and 144 hs after the isolation. N = 3, Mean + SE, n > 100, \*\**p* < 0.01, \*\*\**p* < 0.001.

### Baculoviral transduction of PIEZO1-EGFP into myotubes generated from isolated MuSCs

Non-dividing cells are generally considered resistant to virus-mediated gene transduction (Arnett et al., 2014; Lee et al., 2007; Valeri et al., 2024). Myotubes formed and terminally differentiated by the fusion of myoblasts are one example of non-dividing cells. To determine whether PIEZO1 can be expressed in myotubes using our system, we induced differentiation by culturing isolated MuSCs, followed by low-serum culture to drive myotube formation. As shown in Fig. 4A, baculovirus expressing *Piezo1-egfp* was infected either just before (shown as 6 day) or after (shown as 8 day) induction of myotube formation. Under both conditions, cells that were positive for both Myosin Heavy Chain (MyHC) and EGFP were observed (Fig. 4B, red arrowheads): however, a substantial number of EGFP-positive/ MyHC-negative cells were observed in cells infected following the induction of myotube formation (Fig. 4B, white arrowheads), which is consistent with the findings of a previous study using primary trout salmon myoblasts (Jackson et al., 2013). The results suggest that baculovirus infection before the induction of myotube formation is suitable for the efficient expression of PIEZO1 in myotubes.

**Figure 4.**
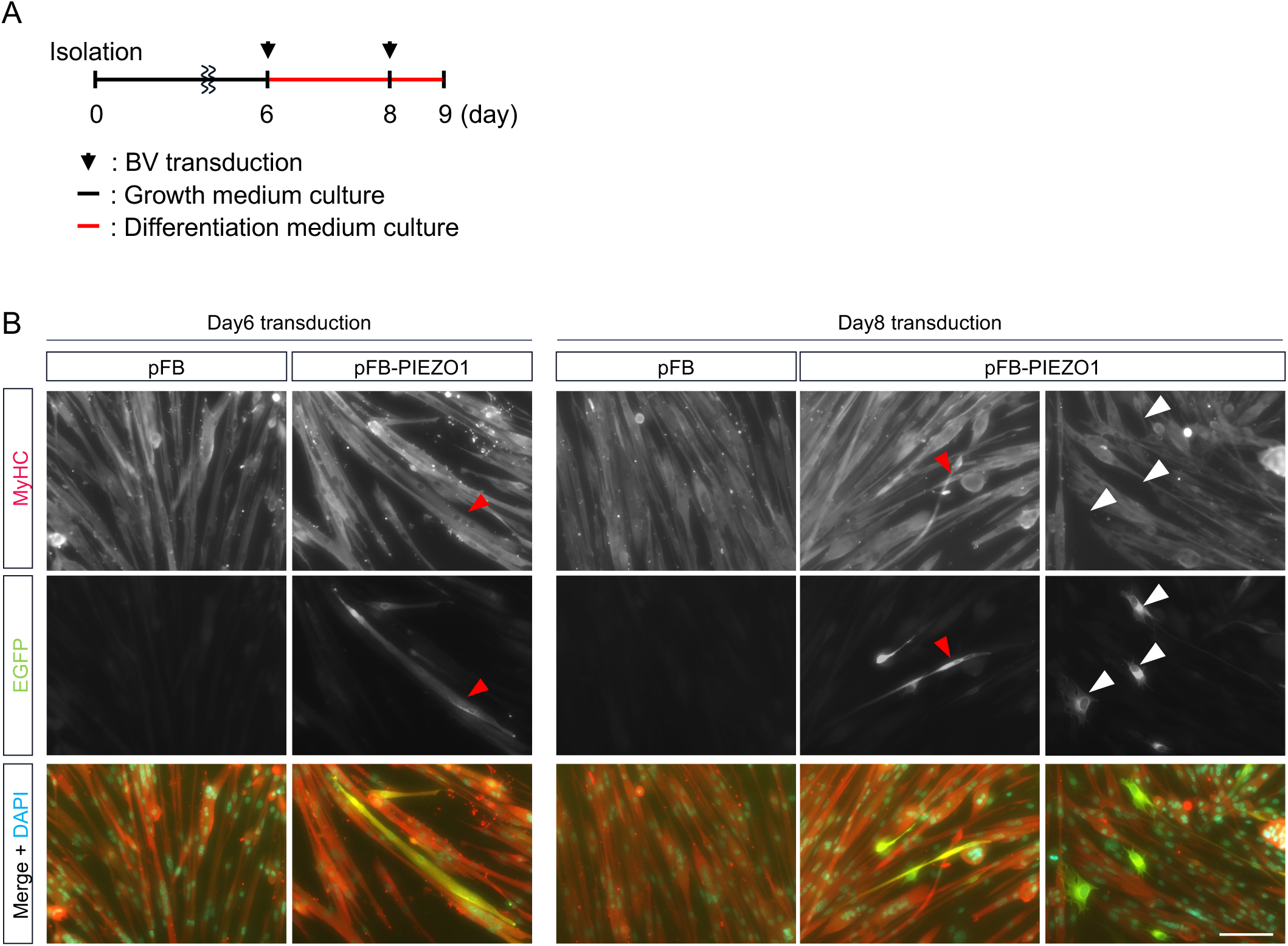
Baculovirus-mediated PIEZO1 expression in differentiating muscle satellite cells. (A) Time course for differentiation of isolated MuSCs and the baculovirus-mediated gene transfer. (B) Immunofluorescent staining of PIEZO1 in differentiating MuSCs transduced with the recombinant baculovirus before (Day 6) or after (Day 8) induction of differentiation. Blue, green, and red represent nuclei, PIEZO1, and MyHC, respectively. Red arrowheads: MyHC and EGFP-positive cells, white arrowheads: MyHC-negative and EGFP-positive cells. Scale bar: 100 μm.

### PIEZO1 expressed via baculovirus in isolated MuSCs retains the ion channel activity

To evaluate the ion channel properties of the expressed PIEZO1-EGFP, MuSCs were evaluated 24 h after isolation, when *Piezo1* was most efficiently introduced by the baculovirus vectors. An immunoblot analysis revealed that the full-length PIEZO1-EGFP protein was expressed only in cells infected with pFB-PIEZO1 (Fig. 5A). Consistent with the results obtained in C2C12 cells, immunofluorescence staining indicated that PIEZO1-EGFP was localized to the plasma membrane (Fig. 5B). Ca²⁺ imaging using Fura-2 AM was conducted in the MuSCs isolated from *Piezo1*-cKO or wild-type mice, and Yoda1-induced Ca²⁺ influx was assessed in each group. Although infection with pFB-PIEZO1virus did not further enhance Yoda1-induced Ca²⁺ influx (Δratio) in wild-type MuSCs, the complete loss of responsiveness to Yoda1 in MuSCs isolated from *Piezo1*-cKO mice was fully restored by pFB-PIEZO1 (Fig. 5C). These results indicate that our baculovirus-based gene transfer system enables the functional expression of PIEZO1 ion channels at the plasma membrane of isolated MuSCs. Moreover, the system is capable of restoring PIEZO1 protein expression in *Piezo1*-deficient MuSCs to comparable levels as that in wild-type cells.

**Figure 5.**
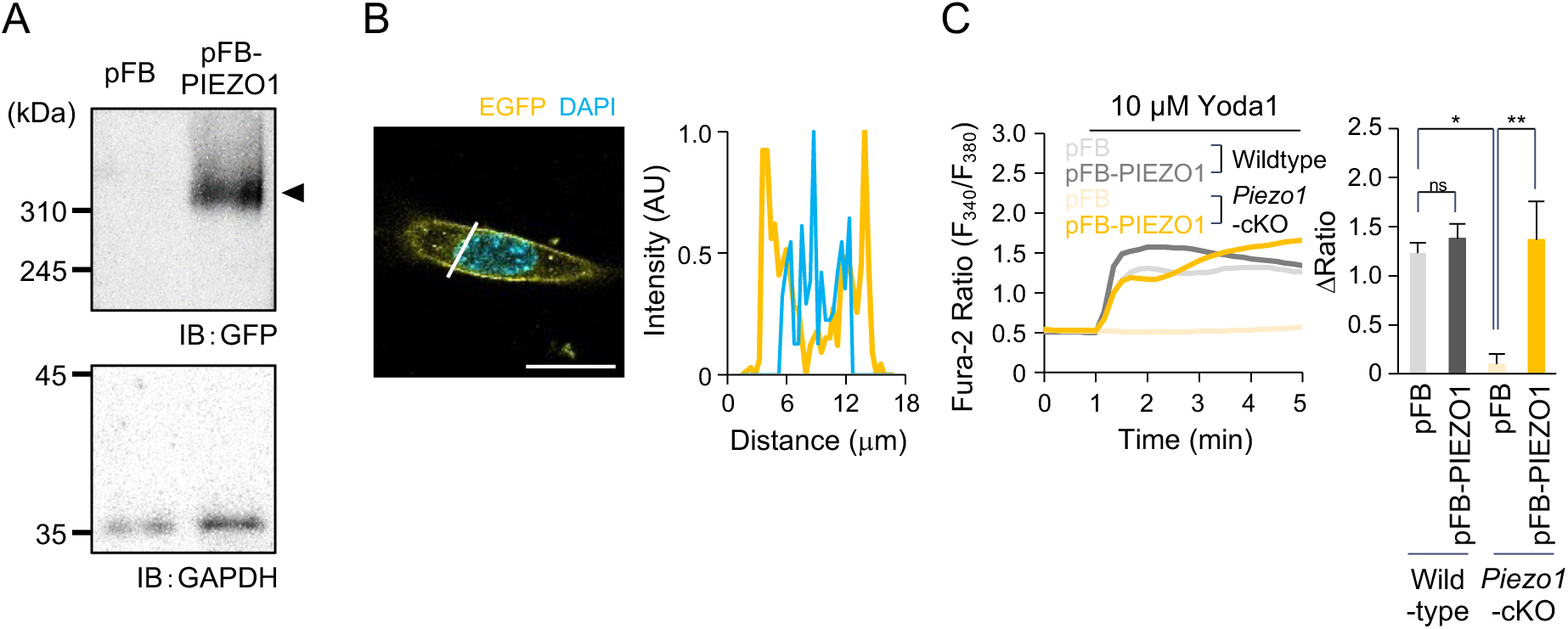
Functional characterization of PIEZO1-EGFP as an ion channel expressed using a baculovirus system in isolated MuSCs. (A) Immunoblot analyses of PIEZO1 protein expression. Isolated MuSCs were transduced with recombinant baculovirus 24 h after the isolation. PIEZO1 and GAPDH were detected using an anti-GFP antibody and an anti-GAPDH antibody, respectively. GAPDH was used as loading control. (B) Fluorescence intensity profiles of PIEZO1-EGFP in the isolated MuSCs transduced with recombinant baculovirus. The intensity profile along the white line was shown in the upper panel. PIEZO1-GFP was detected using anti-GFP antibody (yellow). DAPI (blue) indicated the nuclei. Scale bar: 20μm. (C) Fura-2 ratiometric measurements (F340/F380) of Yoda1-induced Ca^2+^ influx in MuSCs transduced with the recombinant baculovirus (left). MuSCs were isolated from wild type (WT, N = 4) or MuSCs-specific *Piezo1*-deficient (*Piezo1*-cKO, N = 3) mice and 99 cells were analyzed in each assay. Changes in Fura-2 ratio (1′Ratio) were quantified in right panel. Mean + SE, \**p* < 0.05, \*\**p* < 0.01; n.s., not significant.

### PIEZO1 expressed via baculovirus contributes to muscle satellite cell proliferation in a manner comparable to endogenous PIEZO1

The function of ectopically expressed PIEZO1 protein was examined during cell division using isolated MuSCs. At approximately 40 h post-isolation, when the first cell division occurs, PIEZO1-EGFP was co-localized with Aurora kinase (AURORA), which accumulates at the midbody during mitosis (Fig. 6A, left panel). PIEZO1-EGFP also co-localized with RHOA, which is involved in PIEZO1-dependent signaling pathways (Fig. 6A, right panel). This suggests that the exogenously expressed PIEZO1-EGFP may be functional at the midbody during cell division.

**Figure 6.**
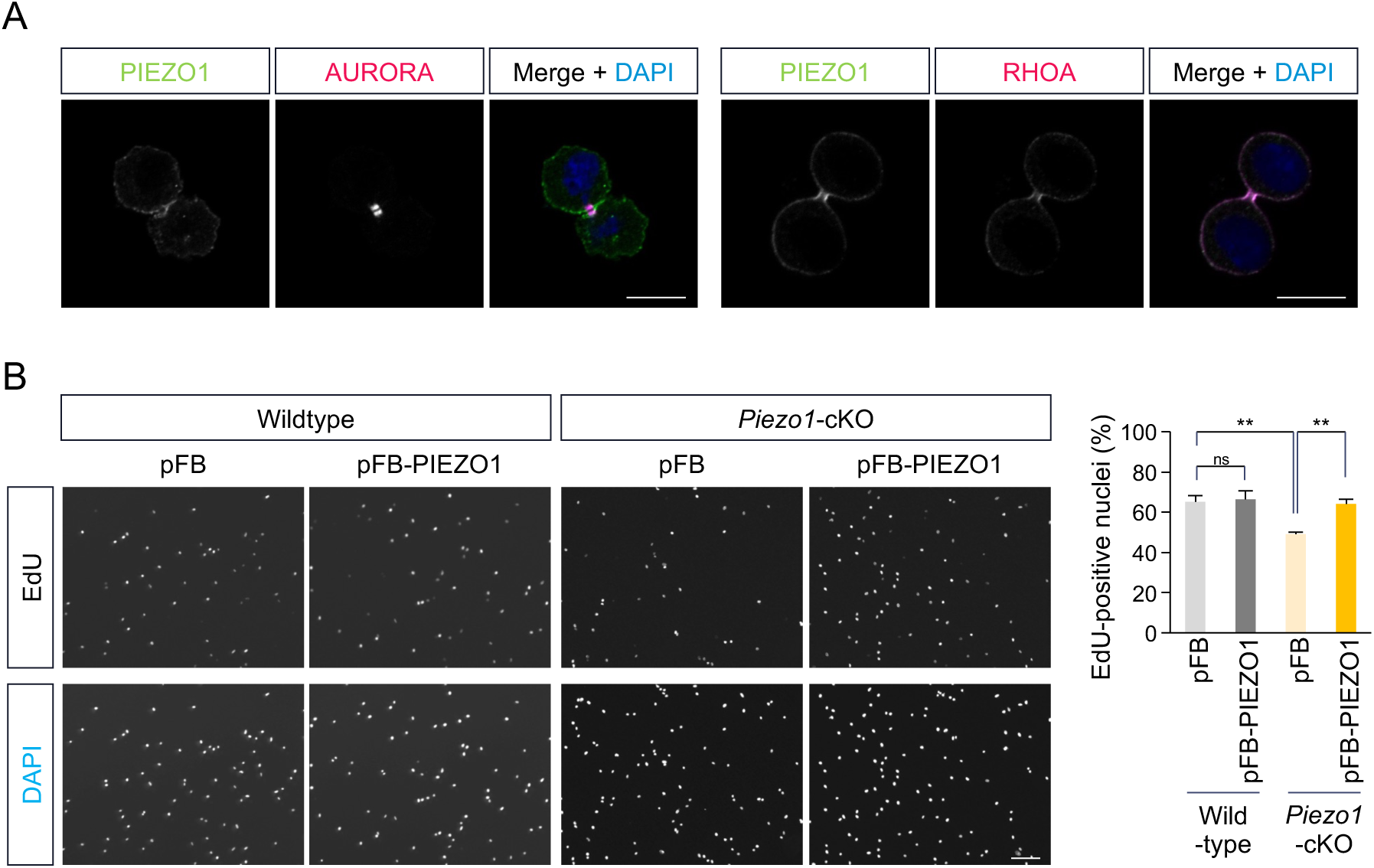
Functional evaluation of baculovirus-expressed PIEZO1-EGFP in proliferating MuSCs. (A) The fluorescence immunostaining analysis of PIEZO1-EGFP localization during cytokinesis. The isolated MuSCs were transduced with recombinant baculovirus 24 h after the isolation. AURORA and RHOA were detected as cytokinetic components using specific antibodies. Scale bar: 10 μm. (B) Fluorescence images of 5-ethynyl-20-deoxyuridine (EdU) incorporation assay in MuSCs transduced with the recombinant baculovirus 24 h after the isolation (left). MuSCs were isolated from wild type (WT, N = 3) or MuSC-specific *Piezo1*-deficient (*Piezo1*-cKO, N = 5) mice and > 400 DAPI-positive cells were analyzed in each assay. Percentages of EdU- and DAPI-positive cells were quantified in right panel. The rate of DNA biosynthesis was evaluated by measuring EdU incorporation 72 h after isolation. Scale bar: 100 μm. Mean + SE, \*\**p* < 0.01; n.s., not significant.

Finally, we determined whether exogenously expressed PIEZO1 could restore the proliferative capacity of *Piezo1*-cKO MuSCs. Consistent with our previous findings (Hirano et al., 2023), EdU incorporation assays revealed a lower percentage of EdU-positive cells in *Piezo1*-cKO MuSCs compared with the wild-type cells. Notably, infection with pFB-PIEZO1 restored the EdU-positive population in *Piezo1*-cKO MuSCs to that of wild-type cells (Fig. 6B). These results indicate that our baculovirus vector system enables the functional expression of the large ion channel PIEZO1 in isolated MuSCs, and the exogenously expressed PIEZO1 is capable of regulating various processes in MuSCs, such as cell proliferation.

## Discussion

During our previous studies of the PIEZO1 ion channel, we recognized the necessity of establishing a system to ectopically express *Piezo1* in isolated MuSCs: however, standard lipofection methods failed to express *Piezo1* likely due to its large gene size, even in C2C12 myoblast cell line (Supplementary Fig. 1). To overcome this obstacle, we constructed a VSVG-pseudotyped baculovirus expressing EGFP-tagged PIEZO1 under the control of the CMV promoter, optimized the infection conditions, and successfully achieved efficient transfer and expression of the *Piezo1* gene in C2C12 cells (Fig. 1). The optimized infection conditions were also applicable to MuSCs isolated from mice, thus allowing the functional expression of PIEZO1 as an ion channel (Figs. 2-5). PIEZO1 expressed using this system fully restored the proliferation defect observed in *Piezo1*- deficient MuSCs (Fig. 6). Although a previous study reported the application of a baculovirus expression vector to MuSCs isolated from trout salmon, this is the first report of exogenous gene expression in mammalian MuSCs using a baculovirus system. The results provide a valuable tool for examining the function of PIEZO1 and other large genes in MuSCs, as well as in other primary cell types.

To maximize the transduction efficiency of *Piezo1*, we established optimal infection conditions for baculovirus-mediated gene transfer. Previous studies have indicated that baculovirus infection under NaHCO₃-free medium or PBS at 25–27°C, enhances gene transfer efficiency in mammalian cells (Hsu et al., 2004; Shen et al., 2007). Consistent with these findings, we found that infection at 27°C in PBS resulted in higher PIEZO1 expression in C2C12 cells compared with infection at 37°C in DMEM (Fig. 1C). Baculovirus are assumed to enter cells through endocytosis, with the low pH of the endosome triggering fusion between the viral envelope and the endosomal membrane, thus enabling release of the viral nucleocapsid into the cytosol (Blissard and Theilmann, 2018). Moreover, low pH can promote direct fusion of the viral envelope with the plasma membrane, followed by immediate cytosolic release of the nucleocapsid (Dong et al., 2010). These findings suggest that low pH may enhance gene transfer efficiency. Therefore, we measured the pH of the medium before and after infection under four conditions: however, no correlation was evident between extracellular pH and the level of PIEZO1 protein expression (Supplementary Fig. 5). Future studies such as monitoring intracellular pH dynamics may identify the mechanism by which the established conditions enhanced PIEZO1 expression (Chin et al., 2021; Rennick et al., 2022). Such mechanistic insights may also facilitate broader application of this baculovirus system to the expression of other genes or primary cell types.

The efficiency of gene expression mediated by viral vectors is influenced by the physiological states of the target cells. Although lentiviral vectors are widely used for gene transfer into isolated stem cells, their transduction efficiency is lower in quiescent (G_0_ phase) cells compared with cells entering the cell cycle (Valeri et al., 2024). Similarly, baculovirus-mediated gene transfer was found to be less efficient in quiescent chondrocytes (Lee et al., 2007). These observations are consistent with our results (Fig. 4), in which MuSCs infected 24 h after isolation (i.e., activated MuSCs) showed a higher PIEZO1 expression efficiency compared with those infected 0.5 h following isolation (i.e., quiescent MuSCs). One possible explanation for this observation is that activated MuSCs may be more susceptible to baculovirus transduction. A series of cellular processes is involved in baculovirus transduction, including attachment to the host cell membrane, internalization, and membrane fusion with the endosome membrane (Matreyek and Engelman, 2013; Nowakowski et al., 2016). Several proteins have been proposed as putative host factors for baculovirus entry into mammalian cells, including the LDL receptor (*Ldlr*) and Niemann-Pick disease type C (*Npc*) (Finkelshtein et al., 2013; Gretch et al., 1991; Huang et al., 2024). A previous transcriptomic profiling study of changes in gene expression during MuSC activation and differentiation revealed that *Npc2* expression was upregulated following activation (Supplementary Fig.6) (Harada et al., 2018). This supports our hypothesis that viral receptor expression may contribute to the higher efficiency of baculovirus-mediated transduction in activated MuSCs. Further studies into the relationship between the expression levels of these putative receptors and viral transduction efficiency were warranted to further improve gene transfer strategies using baculovirus vectors in isolated MuSCs.

Because baculovirus exhibits a large packaging capacity and a non-integrative nature in mammalian cells (Airenne et al., 2013), our system represents an effective tool for future studies of MuSCs. First, this system may be applied to the transduction of PIEZO1 proteins fused with various fluorescent proteins or peptide tags into isolated MuSCs, which may provide further insight into PIEZO1 function in MuSCs. We succeeded in functionally expressing PIEZO1 containing a 3xFLAG tag as an active ion channel in C2C12 cells (Supplementary Fig. 7), which provides a framework for identifying PIEZO1-interacting proteins in MuSCs through co-immunoprecipitation approaches. A major advantage of our system is its ability to efficiently introduce genes into inactive (quiescent) MuSCs. Isolated MuSCs maintained in a quiescent state are increasingly recognized as a promising platform for *ex vivo* gene therapy in genetic muscle disorders beyond their value as a model for cell biological studies. As shown in Fig. 3C, our baculovirus vector successfully mediated the expression of the large PIEZO1 protein in 24% of MuSCs immediately after isolation, which yielded a high gene transfer efficiency into a quiescent population. Therefore, further development and refinement of this system may contribute not only to studies of MuSC biology but also to the advancement of stem cell-based therapies for a wide range of muscular diseases.

## Methods

### Mice

Animal care, ethical use, and protocols were approved by the Animal Care Use and Review Committee of the University of Shizuoka. Male C57BL/6J mice (6-8 weeks of age) were purchased from Japan SLC (Hamamatsu, Japan). Muscle satellite cell (MuSC)-specific *Piezo1*-deficient (*Piezo1*-cKO) mice were generated according to our previous report (Hirano et al., 2023). Briefly, *Piezo1*-floxed mice were mated with *Pax7^CreERT2^*^/+^ transgenic mice (Lepper and Fan, 2010). To induce *Cre* recombinase expression, *Piezo1^flox/flox^*; *Pax7^CreERT2^*^/+^ mice were treated with tamoxifen (TMX; Sigma-Aldrich). TMX was dissolved in corn oil at a concentration of 20 mg ml^−1^ and injected into mice intraperitoneally at 40 μg g^−1^ of body weight for five consecutive days and before isolation of skeletal muscle.

### MuSC isolation using FACS

MuSCs from limb muscles were isolated as previously described (Hirano et al., 2025; Hirano et al., 2023). Briefly, skeletal muscle samples obtained from the forelimbs of mice were subjected to collagenase treatment using 0.2% collagenase type II (Worthington). Mononuclear cells were incubated with fluorescently labeled antibodies, such as anti-mouse Ly-6A/E (Sca-1) (#122508; BioLegend), anti-mouse CD45 (#103106; BioLegend), anti-mouse CD31 (#102508; BioLegend), and anti-mouse CD106 antibodies (#105718; BioLegend) at 4°C for at least 30 min. These cells were resuspended in PBS containing 2% FBS and then subjected to cell sorting to collect CD106-positive cells using BD FACS Aria II or SONY MA 900.

### Cell culture

C2C12 were purchased from ATCC and the *Piezo1*-deleted cell line was previously established using CRISPR/Cas9 system (Tsuchiya et al., 2018). These cells were grown in 10% FBS/DMEM (GM). MuSCs isolated from mice were cultured in growth medium (SC-GM; DMEM supplemented with 30% fetal bovine serum (Sigma-Aldrich), 1% chick embryo extract (US Biological), 10 ng ml^−1^ basic fibroblast growth factor (ORIENTAL YEAST Co., Ltd.), and 1% penicillin-streptomycin (FUJIFILM Wako Pure Chemical Corporation)) on culture dishes coated with Matrigel (Corning). Cells were cultured in a 37°C incubator with 5% CO_2_. For inducing myotube formation, C2C12 cells semi-confluently grown in 10% FBS/DMEM were maintained in differentiation medium (DM; DMEM containing 2% horse serum (Gibco)) for the indicated periods. The culture medium was replaced with fresh DM every other day. Isolated MuSCs grown in SC-GM for 6 days were maintained in differentiation medium (SC-DM; DMEM containing 5% horse serum and 1% penicillin-streptomycin) for 2 days.

### Baculovirus production

Baculovirus that expresses vesicular stomatitis virus G-protein (VSVG) on the virus envelope was produced using the Bac-to-Bac system (Life Technologies) as described previously (Murayama et al., 2022). Mouse *Piezo1* cDNA was cloned into the modified donor vector (pFastBac1-VSVG-CMV-WPRE). Baculovirus was produced in Sf9 cells according to the manufacturer’s instructions (Life Technologies). Titers of baculovirus, immunologically determined using an anti-gp64 antibody (AcV5, Santa Cruz Biotechnology), were approximately 1 × 10^8^ pfu ml^−1^ for all viruses. The baculovirus solutions were stored in the dark at 4°C. For baculovirus infection, C2C12 cells and isolated MuSCs were cultured in baculovirus solutions mixed with DMEM or PBS at multiplicities of infection (MOI) of 400 and 100, respectively, for 4 h at 27°C or 37°C.

### Immunoblot analysis

Cells were washed with PBS and lysed in lysis buffer (10 mM Tris-HCl (pH 7.4), 1% Triton X-100, 0.1% sodium dodecyl sulfate (SDS), 1% sodium deoxycholate) containing 1% protease inhibitors (Nacalai Tesque). The lysates were centrifuged at 14,000 × g for 10 min at 4°C. Then 40 μl of supernatant was mixed with 10 μl of SDS sampling buffer 1 (50 mM Tris-HCl (pH 8.0), 50% sucrose, 1% SDS, 5 mM ethylenediamine tetraacetic acid (EDTA), 0.4% bromophenol blue) and incubated for 10 min at 50°C, followed by mixing with 50 μl of SDS sampling buffer 2 (10 mM Tris-HCl (pH 8.0), 10% sucrose, 0.2% SDS, 1 mM EDTA, 0.08% bromophenol blue, 60% urea). The samples were electrophoresed on 10% SDS-polyacrylamide gel, and then were blotted on polyvinylidene fluoride (PVDF) membranes using Trans-Blot SD Semi-Dry Electrophoretic Transfer Cells (Bio-Rad). Immunoblot analysis was performed using rabbit anti-GFP antibody (#598; 1:500; MBL), rabbit anti-α Tubulin antibody (1:1,000, Cell Signaling, #2144), and mouse anti-GAPDH antibody (#CB1001; 1:1,000; Sigma-Aldrich). Bound antibodies were detected using horseradish peroxidase-conjugated anti-rabbit immunoglobulin (Ig) G antibody, anti-mouse IgG antibody, or anti-rat IgG using Super Signal West Pico (Thermo Scientific) and Ez-Capture MG (Atto).

### Immunofluorescent analysis

Cells were fixed with 4% paraformaldehyde (PFA)/PBS, permeabilized in 0.5% Triton X-100/PBS, blocked in 1% BSA/PBS, and probed with antibodies. For RhoA detection during cytokinesis, isolated MuSCs were fixed with 10% trichloroacetic acid on ice for 15 min. They were then rinsed twice with PBS containing 30 mM glycine (G-PBS) and treated with 0.2% Triton X-100 in G-PBS for 5 min for permeabilization (Hirano et al., 2023). Cells were stained with anti-GFP (AB0020-200; 1:200; SICGEN), anti-RHOA (sc-418; 1:100; Santa Cruz), and anti-Phospho-AURORA antibodies (2914S; 1:200; Cell Signaling Technology) with DAPI (Dojindo), and observed using a confocal microscope (LSM800, Zeiss) with a 63× objective lens. To assess myotube formation, cells were stained with anti-MyHC antibody (14-6503-82; 1:200; eBioscience) and DAPI (Dojindo), and observed using an epifluorescence microscope (Axio-Observer.Z1, Zeiss) with a 10 × objective lens. To evaluate the activated and differentiated states of MuSCs, cells were stained with anti-PAX7 (Pax7-c; 1:100; DSHB) or anti-MYOD (sc-304; 1:500; Santa Cruz) antibody and DAPI (Dojindo), and observed using an epifluorescence microscope (Axio-Observer.Z1, Zeiss) with a 10× objective lens. The number of DAPI-stained nuclei was counted in each microscopic field using ImageJ software.

### Ca^2+^ imaging

C2C12 cells or isolated MuSCs were seeded onto glass-bottomed dishes coated with poly-L-lysine or Matrigel, respectively. Cells were loaded with 5 μM Fura-2 AM (Dojindo Laboratories) in culture medium at 37 °C for 60 min. Time-lapse imaging was performed at 10-second intervals. The base composition of HBS was 140 mM NaCl, 5 mM KCl, 2 mM MgCl_2_, 2 mM CaCl_2_, 10 mM glucose, and 10 mM HEPES (pH = 7.4 adjusted with NaOH). Ratiometric imaging (F340/F380) of Fura-2 fluorescence was conducted using Physiology software (Zeiss). Yoda1-induced Ca^2+^ influx was measured as the difference in the Fura-2 ratio (Δratio) between its maximum value and that at 1 min from the initiation of imaging.

### Evaluation of MuSC proliferation

Isolated MuSCs were seeded onto 24-well plates coated with Matrigel (Corning), and were incubated with GM containing 5 mM EdU (Life technologies) for 3 h. For the detection of the incorporated EdU, a click chemical reaction was performed using a Click-iT EdU Imaging Kit (Life Technologies) after primary and secondary antibody staining, according to the manufacturer’s instructions.

### Measurement of pH of the culture medium

The baculovirus medium (Sf-900II™ SFM, Gibco) was mixed with DMEM or PBS at a 1:4 ratio, and the pH was measured immediately (shown as 0 h) using a pH meter (HORIBA). To evaluate pH changes in the mixed medium after cell culture, 1 × 10^5^ C2C12 cells were seeded in a 10-cm dish and cultured overnight. The medium was the replaced with the mixed medium and incubated at either 27°C or 37°C for 4 h. The pH of the culture supernatant was measured after centrifugation (shown as 4 h).

### Statistical analysis

Statistical analyses were performed using Microsoft Excel, R version 3.6.3 (2020-02-29). The statistical significance of the differences between the mean values was analyzed using a non-paired t-test (two-sided). Multiple comparisons were performed using Tukey’s test followed by analysis of variance (ANOVA). P values of \**P* < 0.05, \*\**P* < 0.01, and \*\*\**P* < 0.001 were considered statistically significant.

## Acknowledgments

We thank members of the Hara laboratory for their scientific contributions. The mouse strain (*Piezo1^tm1c(KOMP)Wtsi^*) used for this research project was created by the Mouse Biology Program (www.mousebiology.org) at the University of California Davis, generated by the trans-NIH Knock-Out Mouse Project (KOMP) and obtained from the MMRRC (www.mmrrc.org). NIH grants to Velocigene at Regeneron Inc (U01HG004085) and the CSD Consortium (U01HG004080) funded the generation of gene-targeted vectors and ES cells for over 8500 genes for the KOMP Program. This study was supported by the Grant-in-Aid for Scientific Research KAKENHI (22H03484, 25K02994); Grant-in-Aid for Transformative Research Areas B (23H03856); Intramural Research Grant (5-6) for Neurological and Psychiatric Disorder of NCNP; grants from Takeda Science Foundation, ONO Medical Research Foundation, Uehara memorial foundation, Chugai Foundation for Innovative Drug Discovery Science (SRG2022), and the Asahi Glass Foundation (to Y.H.); and the Grant-in-Aid for Scientific Research KAKENHI (22K20636, 23K14184), UBE Foundation, ONO Medical Research Foundation (to A.Mu.).

## Author contributions

Conceptualization: Y.H.; Methodology: K.H, M.T., Y.On., and T.M.; Investigation: A.Mu., A.Ma., T.H.; Supervision: A.Mu., Y.H.; Writing—original draft: A.Mu.; Writing—review and editing: A.Mu., Y.H.

## Data and material availability

All data needed to evaluate the conclusions in the paper are present in the paper.

---

**Supplementary Figure 1.**
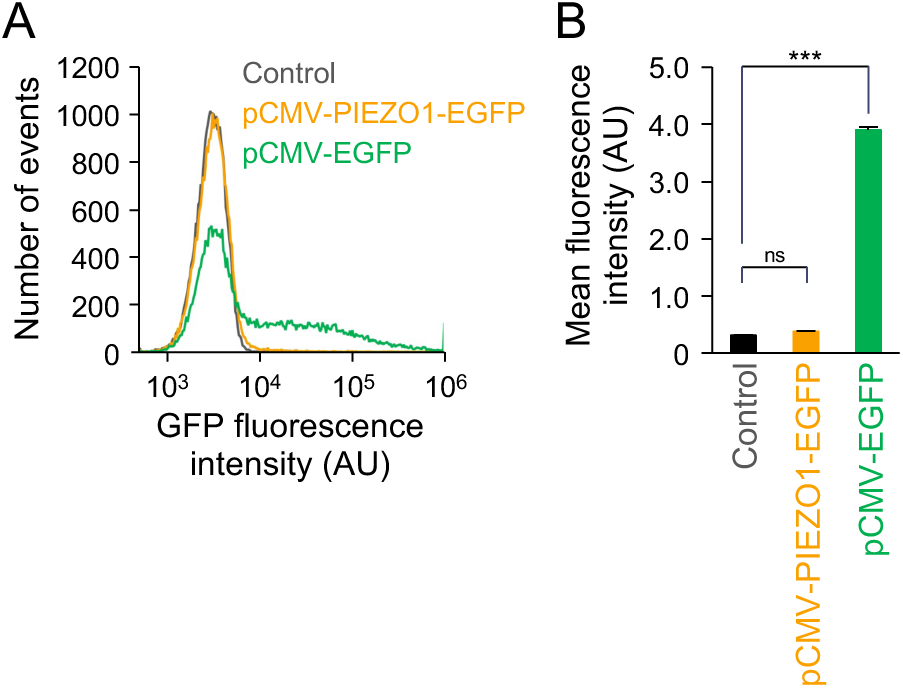
Low transfection efficiency of a plasmid encoding PIEZO1-EGFP into C2C12 cells using a lipofection reagent. (A) Flow cytometry analyses of GFP fluorescence intensity in C2C12 cells transfected with pCMV-EGFP plasmid (green) or pCMV-PIEZO1-EGFP plasmid (yellow) using Lipofectamine 2000. (B) The mean fluorescence intensity among the cells analyzed by the flow cytometry. N = 3, n = 30,000 cells, Mean + SE, \*\*\**p* < 0.001; n.s., not significant.

**Supplementary Figure 2.**
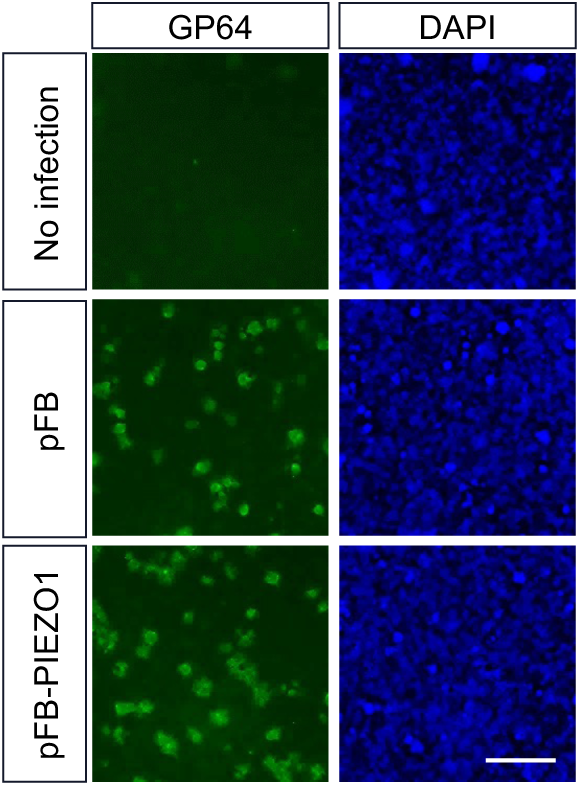
Generation of the recombinant baculovirus. Top panels: uninfected Sf9 cells, middle panels: Sf9 cells infected with pFB baculovirus, bottom panels: Sf9 cells infected with pFB-PIEZO1 baculovirus. GP64 or DAPI represented the baculovirus envelope protein or nuclei, respectively. Scale bar: 500 μm.

**Supplementary Figure 3.**
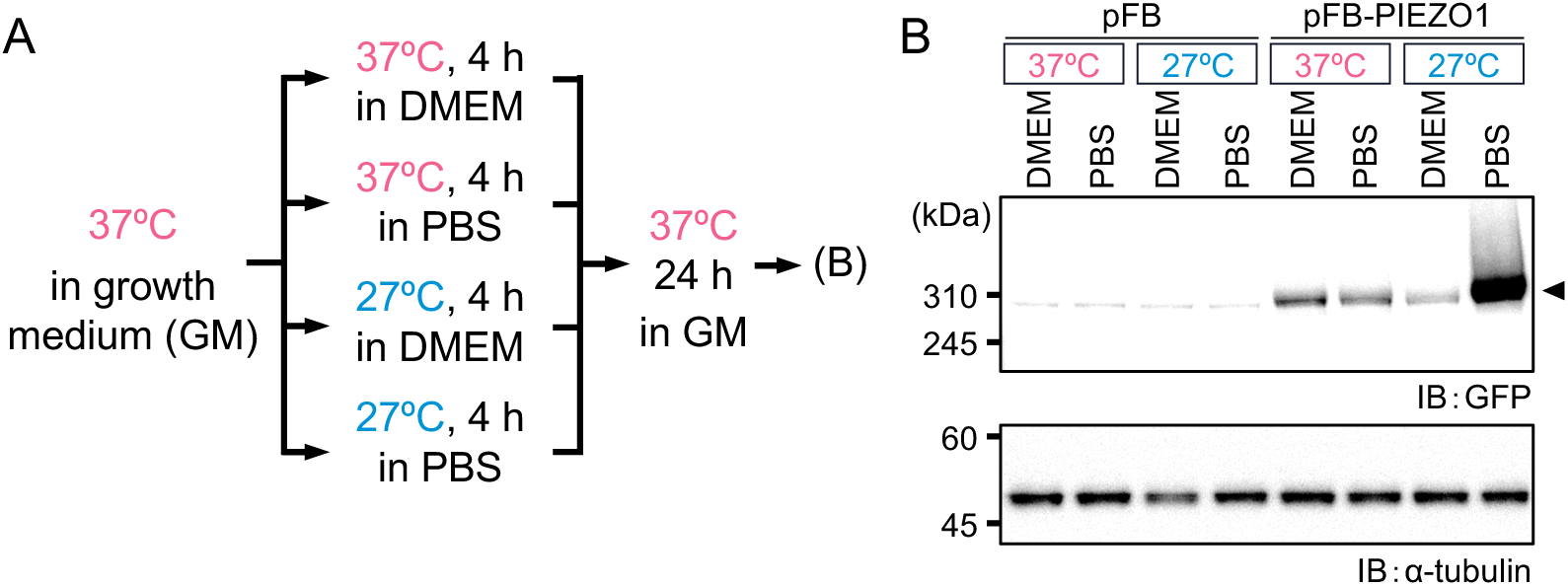
Effects of temperature and pH in culture medium on viral transduction. (A) Baculovirus transduction into C2C12 cells under indicated culture conditions. (B) Immunoblot analyses of PIEZO1 protein expression. C2C12 cells were transduced with recombinant baculovirus under the indicated culture conditions. The amount of PIEZO1 and α-tubulin were detected using an anti-GFP antibody and anti-α-tubulin antibody, respectively. α-tubulin was used as loading control.

**Supplementary Figure 4.**
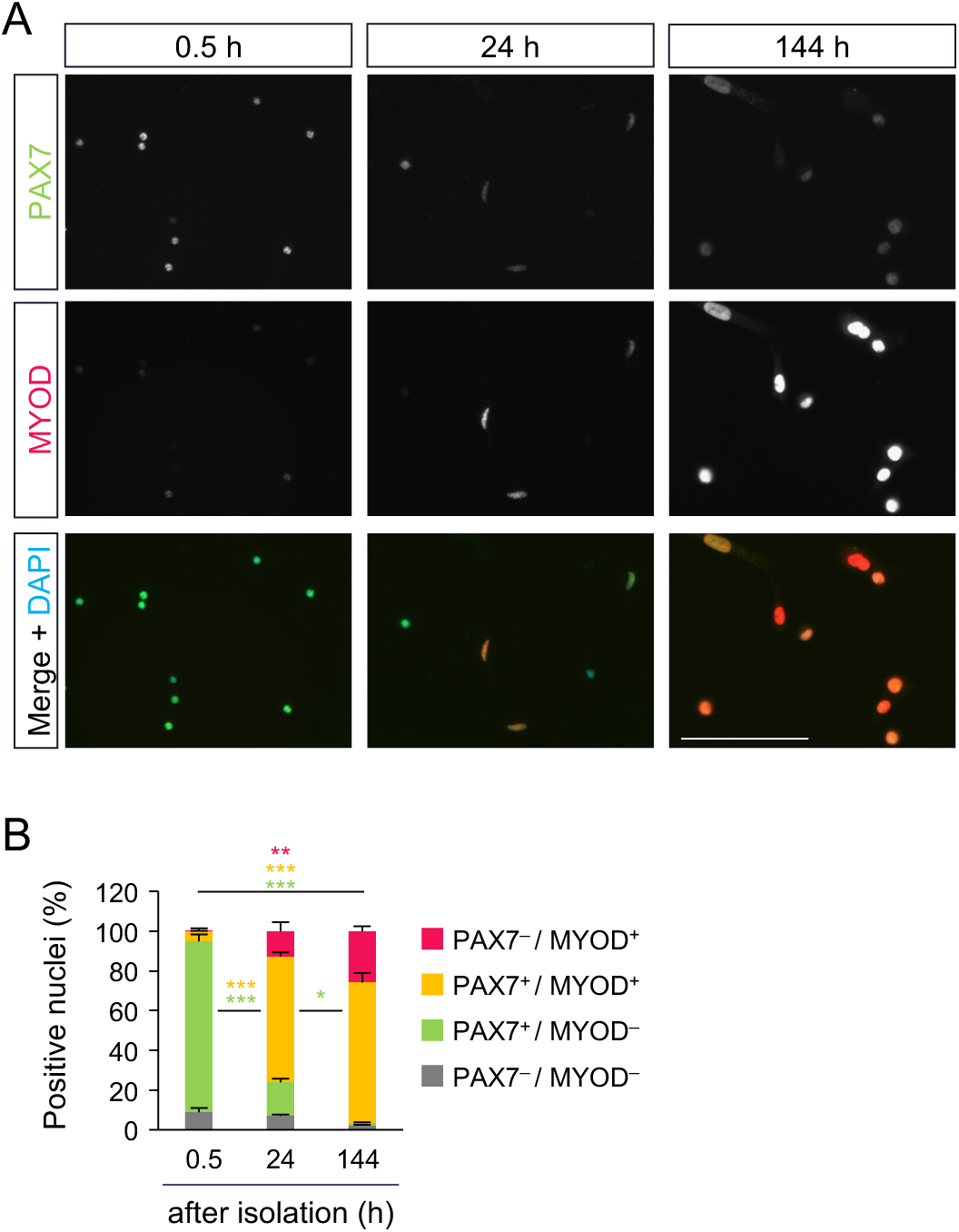
Evaluation of activated/differentiated states in isolated MuSCs before baculoviral transduction. Immunofluorescence analyses of MYOD (red) and PAX7 (green) in muscle satellite cells at the indicated time points after isolation (A). The percentage of MYOD and/or PAX7-expressing cells were quantified (B). DAPI (blue) indicated the nuclei. Scale bar: 100 μm. N = 3, n > 200 cells, Mean + SE, *\*p* < 0.05, \*\**p* < 0.01, \*\*\**p* < 0.001.

**Supplementary Figure 5.**
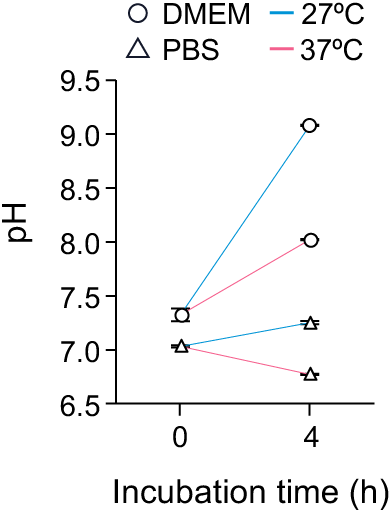
Changes in pH in culture media of C2C12 cells. The pH in culture media was measured after a 4-h-incubation under the indicated conditions. N = 3, Mean + SE.

**Supplementary Figure 6.**
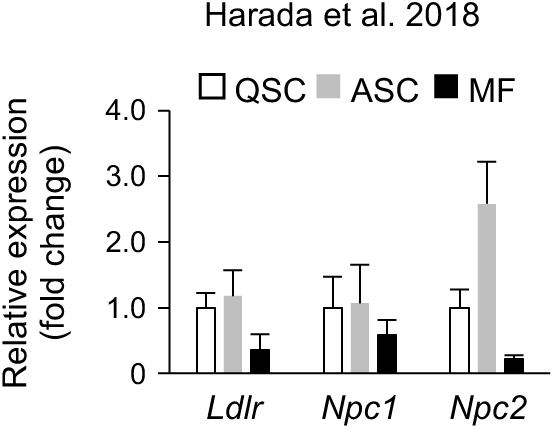
*In silico* analysis of putative baculoviral receptors during MuSC activation/differentiation. Gene expression profiles of putative baculoviral receptors were evaluated in silico using previously-reported RNA-seq data from MuSCs (Harada et al., 2018). QSC: quiescence satellite cell, ASC: activated satellite cell, MF: myofiber. N = 3, Mean + SE.

**Supplementary Figure 7.**
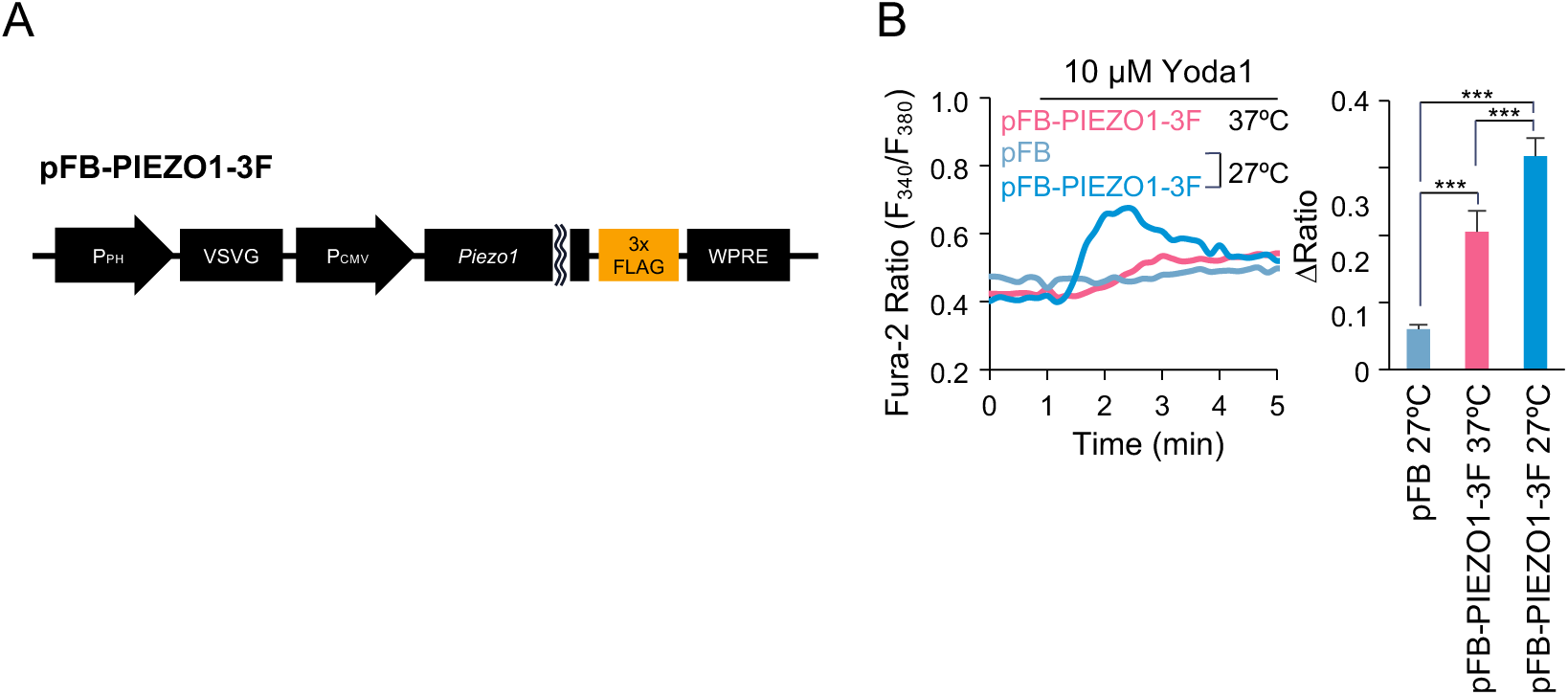
Baculovirus-mediated expression of PIEZO1-3xFLAG in C2C12 cells. (A) Recombinant donor plasmid expressing PIEZO1-3xFLAG (pFB-PIEZO1-3F). PH; polyhedrin, VSVG; vesicular stomatitis virus G-protein, WPRE; woodchuck hepatitis post-transcriptional regulatory element. (B) Fura-2 ratiometric measurements (F340/F380) of Yoda1-induced Ca^2+^ influx in *Piezo1*-deficient C2C12 cells transduced with the recombinant baculovirus under the indicated conditions (left). pFB was used as a control. Changes in Fura-2 ratio (Δratio) were quantified in right panel. n = 99 cells, Mean + SE, \*\*\**p* < 0.001.

